# Sex-specific adaptations to VTA circuits following subchronic stress

**DOI:** 10.1101/2023.08.02.551665

**Authors:** Chloé Bouarab, Megan Wynalda, Brittney V. Thompson, Ambika Khurana, Caitlyn R. Cody, Alexandre Kisner, Abigail M. Polter

## Abstract

Dysregulation of the mesolimbic reward circuitry is implicated in the pathophysiology of stress-related illnesses such as depression and anxiety. These disorders are more frequently diagnosed in females, and sex differences in the response to stress are likely to be one factor that leads to enhanced vulnerability of females. In this study, we use subchronic variable stress (SCVS), a model in which females are uniquely vulnerable to behavioral disturbances, to investigate sexually divergent mechanisms of regulation of the ventral tegmental area by stress. Using slice electrophysiology, we find that female, but not male mice have a reduction in the *ex vivo* firing rate of VTA dopaminergic neurons following SCVS. Surprisingly, both male and female animals show an increase in inhibitory tone onto VTA dopaminergic neurons and an increase in the firing rate of VTA GABAergic neurons. In males, however, this is accompanied by a robust increase in excitatory synaptic tone onto VTA dopamine neurons. This supports a model by which SCVS recruits VTA GABA neurons to inhibit dopaminergic neurons in both male and female mice, but males are protected from diminished functioning of the dopaminergic system by a compensatory upregulation of excitatory synapses.

## Introduction

Stressful experiences have long been associated with changes in the function of the brain’s reward circuitry and reward-related behavior^1^. Stress-induced adaptations in the dopaminergic neurons of the ventral tegmental area (VTA), a major hub of this reward circuitry, are critical to post-stress behavioral sequelae such as anhedonia or potentiated drug seeking^2–5^. Dopaminergic neurons respond non-uniformly to a wide range of acute stressors by both immediate changes in activity and longer-duration shifts in plasticity^6–14^. Fewer studies have investigated responses to chronic or repeated stress, and changes in regulation of dopaminergic activity in these models is highly dependent on the modality, duration, and intensity of the stressor^15–21^. However, many of these studies have largely focused on male rodents, leaving questions of the sex specificity of modulation of VTA function by longer-term stressors unanswered.

Males and females have divergent responses to stress at every level from transcriptional to behavioral^22, 23^. In humans, women are more likely to be diagnosed with stress-related disorders such as depression, PTSD, and anxiety disorders^24, 25^. While there are multiple social and biological factors that contribute to this difference, these data suggest that there are sex-linked mechanisms of both vulnerability and resistance to stress that could be exploited to develop more effective sex-specific treatments for stress induced neuropsychiatric disorders. Subchronic variable stress (Figure 1A) is a six day mouse stress paradigm that provides an opportunity to investigate mechanisms by which stress results in sexually divergent behavioral and physiological responses^26, 27^. In this model, females, but not males, exhibit changes in stress coping, reward consumption and approach-avoidance behavior as well as changes on the transcriptional, immunological, and circuit levels ^26, 28–30^. Here, we use this model to investigate how differential effects on the mesolimbic dopamine system may arise in male and female mice. We show that SCVS has both sex-independent and sex-specific effects on the cellular function of the VTA. These studies suggest that alterations in VTA function may be substrates for active mechanisms of both female-specific vulnerability and male-specific resistance.

**Figure 1:**
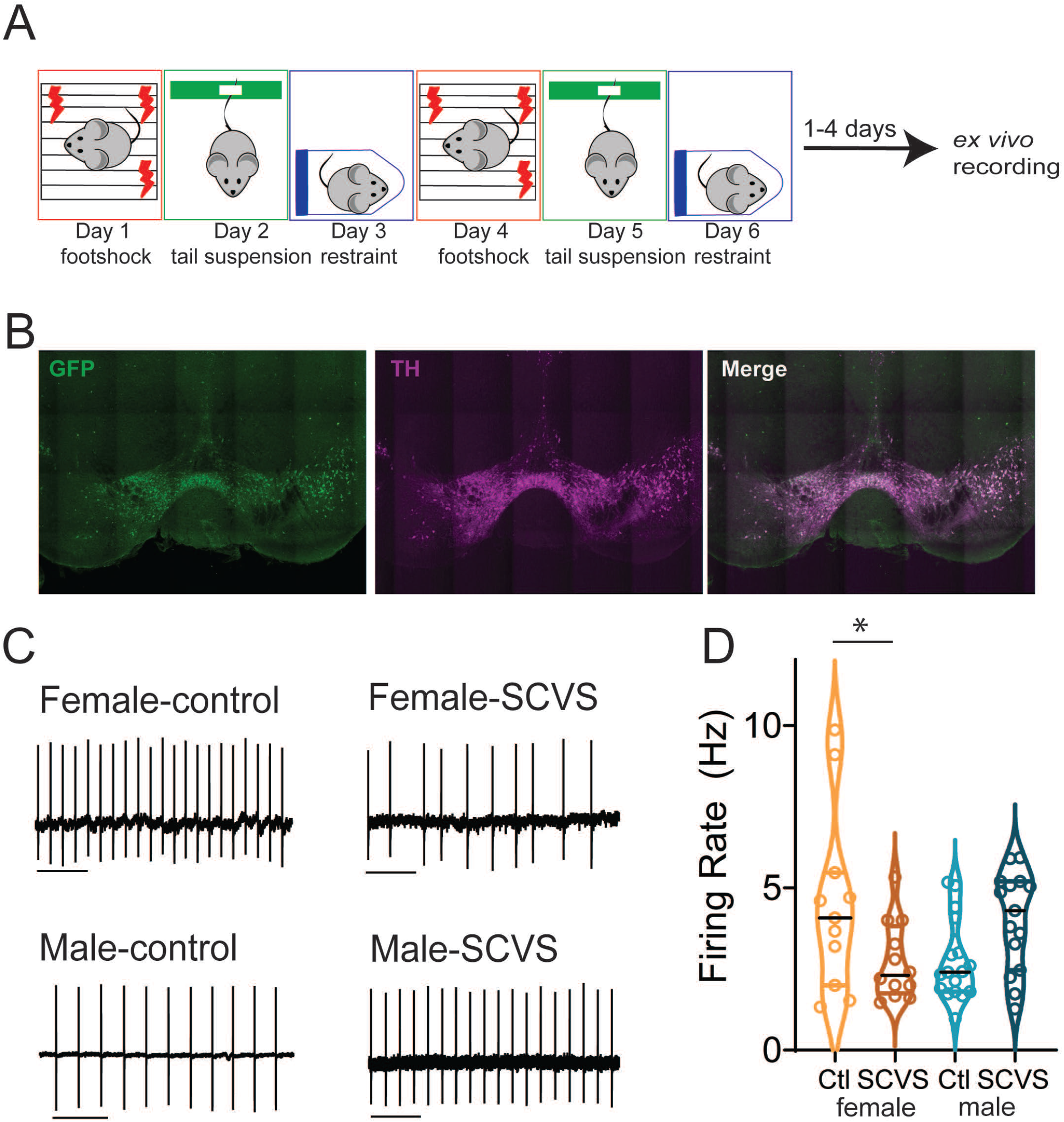
SCVS decreases the firing rate of dopaminergic neurons in female mice. A. Experimental protocol for SCVS and *ex vivo* recording. B. Representative 10 x immunohistochemistry images showing TH expression in GFP+ neurons in the VTA of Pitx3-GFP mice. C. Representative cell-attached traces from dopaminergic neurons. D. Average firing rate of VTA dopaminergic neurons from control and SCVS mice. * p<0.05, 2-way ANOVA followed by Bonferroni test. n=11-17/group. Scale bars=1 s.

## Materials and Methods

### Animals

All animals and experimental protocols were conducted in accordance with National Institutes of Health Guidelines for the Care and Use of Laboratory Animals, and with the approval of the IACUC of The George Washington University. Male and female mice 8-10 weeks of age were used for all studies. For fluorescent identification of VTA dopaminergic neurons, we used Pitx3-GFP mice, which express GFP in dopaminergic neurons (provided by the laboratory of Kevin Wickman, Figure 1B)^31^. For recordings from VTA GABAergic neurons, Vgat-Cre mice (Strain 028862, The Jackson Laboratory)^32^ were crossed with Ai14 tdTomato reporter mice (Strain 007908, The Jackson Laboratory)^33^. Mice were group housed with littermates within ventilated cages in temperature- and humidity-controlled rooms with *ad libitum* access to water and rodent chow on a 12 h light/dark cycle. Stressed and control animals were housed in the same room, but separate cages.

### Immunohistochemistry

Adult Pitx3-GFP mice were deeply anesthetized with ketamine (100 mg/kg) and dexmeditomidine (0.25 mg/kg) and transcardially perfused with 1x Phosphate buffered saline (PBS) followed by 4% paraformaldehyde (PFA). Brains were rapidly dissected and post-fixed at 4°C for 16-24 hours in 4% PFA and then dehydrated in 30% sucrose, and processed for cryo-slicing. 50 micron sections containing the VTA were prepared on a cryostat. VTA sections were stained with a rabbit polyclonal antibody to tyrosine hydroxylase (Sigma-Aldrich, AB152) and a goat anti-rabbit secondary conjugated with Alexa Fluor 594 (Invitrogen, A-11012). Sections were then stained with a chicken anti-GFP antibody (Abcam, 13970) and a goat anti-chicken secondary conjugated with Alexa Fluor 488 (Invitrogen, A-11039). Sections were imaged on a Zeiss Cell Observer Spinning Disk Confocal Microscope.

### Subchronic Variable Stress

The subchronic variable stress (SCVS) protocol was performed as previously described (Figure 1A) ^26, 27^. This paradigm involves daily 1-hour stress sessions of three distinct stressors, footshock (Day 1 and Day 4), tail suspension (Day 2 and 5), and restraint (Day 3 and 6). Footshock was performed in standard electrified fear conditioning box (Coulbourn Instruments) inside a sound attenuation chamber. 100 randomized 0.5 mV footshocks were applied over 60 minutes. Male and female mice were shocked in separate chambers. Control mice were transported with stressed mice and placed in a separate room for an hour. For tail suspension, mice were taped by their tails ∼15 inches above the benchtop for 1 hour. For restraint stress, mice were placed into ventilated 50-mL conical tubes and the tubes were placed inside their home cage for 60 minutes. For both restraint and tail suspension, males and females were stressed in separate sessions and were not exposed to each other. Control mice were transported to the stress room for 60 minutes on tail suspension and restraint days briefly handled, but were not present in the room during stress sessions.

### Acute slice electrophysiology

Electrophysiological recordings were performed between 1 and 4 days following the final stress session. Electrophysiological recordings were performed as previously described ^34, 35^. Pitx3-GFP^31^ or Vgat-Cre:Ai14^32, 33^ mice with fluorescently labeled dopaminergic or GABAergic neurons, respectively, were deeply anesthetized with ketamine (100 mg/kg) and dexmeditomidine (0.25 mg/kg) and perfused transcardially with 34°C N-methyl-D-glucamine (NMDG)-based slicing solution^36^ containing (in mM): 92 NMDG, 20 HEPES, 25 glucose, 30 NaHCO_3_, 1.2 NaH_2_PO_4_, 2.5 KCl, 5 sodium ascorbate, 3 sodium pyruvate, 2 thiourea, 10 MgSO_4_, and 0.5 CaCl_2_. Brains were rapidly dissected and placed in warmed NMDG solution. Horizontal brain slices (220 μm thick) containing the VTA were obtained using a vibratome (Leica VT1200, Leica Biosystems Inc., IL, USA). Immediately after slicing, brain slices were transferred to a holding chamber at 34°C degrees filled with a recovery solution containing (in mM): 92 NaCl, 20 HEPES, 25 glucose, 30 NaHCO_3_, 1.2 NaH_2_PO_4_, 2.5 KCl, 5 sodium ascorbate, 3 sodium pyruvate, 2 thiourea, 1 MgSO_4_, and 2 CaCl_2_. Slices were held at 34°C for one hour, and then at room temperature until use. For electrophysiological recordings, a single slice was transferred to a chamber perfused at a rate of 1.5 to 2.0 ml/min with heated (28-32°C) artificial cerebrospinal fluid (aCSF, in mM: 125 NaCl, 2.5 KCl, 1.25 NaH_2_PO_4_, 1 MgCl_2_ 6H_2_O, 11 glucose, 26 NaHCO_3_, 2.4 CaCl_2_. All solutions were saturated with 95% O_2_ and 5% CO_2_.

In Pitx3-GFP mice, dopaminergic neurons were selected based on GFP fluorescence^31^, presence of I_h_, and location in the lateral half of the VTA. This selection criteria means that our sample is enriched with dopaminergic neurons projecting to the lateral nucleus accumbens, but we cannot rule out the inclusion of dopaminergic cells with other projection targets^37–39^. GABAergic neurons in Vgat-Cre:Ai14 mice were selected based on tdTomato fluorescence and lateral location in the VTA. Whole-cell patch-clamp recordings were performed using a Sutter IPA amplifier (1 kHz low-pass Bessel filter and 10 kHz digitization) using Sutter Patch software (Sutter Instruments). Voltage-clamp recordings were made using glass patch pipettes with resistance 2-4 MOhms, filled with either potassium gluconate (cell-attached and EPSC recordings) or potassium chloride (IPSC recordings) internal solution. Potassium gluconate internal contained (in mM): K-gluconate 117, NaCl 2.8, MgCl2 5, CaCl2 0.2, HEPES 20, Na-ATP 2, Na-GTP 0.3, EGTA 0.6. Potassium chloride internal contained (in mM): 125 KCl, 2.8 NaCl, 2 MgCl2, 2 ATP-Na+, 0.3 GTP-Na+, 0.6 EGTA, and 10 HEPES. Series resistance was monitored throughout voltage clamp recordings and recordings in which the series resistance changed more than 20% and/or exceeded 20 MOhms were not included in the analysis. Membrane potentials in K-gluconate based recordings were not corrected for junction potentials.

Cell attached recordings were performed in aCSF using SutterPatch software in the loose-patch configuration^40^. After a three-minute baseline of stable firing, the firing rate over a 60-second window was measured using SutterPatch’s action potential detection module. Collection of spontaneous and miniature I/EPSCs was performed using SutterPatch software. Cells were voltage-clamped at -70 mV. After a 3-minute stabilization period, a total of 200 spontaneous synaptic events were recorded from each cell. Following collection of spontaneous events, tetrodotoxin (1 µM) was added to the bath. After 10-minute wash-in of tetrodotoxin, a further 200 miniature synaptic events were collected. To isolate GABA_A_R IPSCs, 6,7-dinitroquinoxaline-2,3-dione (DNQX; 10 μM) and strychnine (1 μM) were added to the extracellular solution to block AMPA and glycine receptors respectively. To isolate EPSCs, picrotoxin (100 µM) was added to the aCSF to block GABA_A_ and glycine receptors. Synaptic events were analyzed using the event detection module in SutterPatch with a threshold of 8 pA. IPSCs and EPSCs were detected using the SutterPatch event detection module using a template with a rise of 1400 ms and decay of 6500 ms (IPSCs) or a rise of 1400 ms and a decay of 2000 ms (EPSCs).

### Materials

All salts used for electrophysiology were purchased from Sigma-Aldrich (St. Louis, MO) or Fisher Scientific (Hampton, NH). Pharmacological reagents such as picrotoxin, DNQX, strychnine, and tetrodotoxin were purchased from Tocris Biosciences (Bristol, United Kingdom). Ketamine and dexmedetomidine were purchased from Covetrus (Elizabethtown, PA).

### Statistics

Results are reported in the text as mean ± SEM. For all studies, the n represents the number of animals. Graphs are shown as violin plots or truncated violin plots with medians and quartiles. Data was analyzed using a 2-way ANOVA with sex and stress as factors. If a significant interaction was detected, Bonferroni’s post-hoc test was used to determine significance between stress conditions within each sex. Statistical tests were performed in GraphPad Prism 9.2.

## Results

### SCVS reduces the firing rate of dopaminergic neurons in female mice

SCVS leads to female-specific alterations across a number of behavioral domains, including reductions in active coping behavior^27^, social and general avoidance^26, 28^, and anhedonia^26, 30^. As these behaviors are all profoundly influenced by the mesolimbic dopaminergic system^21, 41–45^, we therefore hypothesized that exposure to SCVS may decrease the functionality of the mesolimbic dopaminergic system in female mice. We subjected male and female pitx3-GFP mice to SCVS (figure 1A-B) and one to four days following the final stress session, we prepared acute VTA slices and performed cell attached recordings to determine the basal firing rate (figure 1C). We found a significant interaction between stress and sex (Figure 1D, two-way ANOVA, F _(1, 51)_ = 10.40, P=0.0022), but no significant effect of stress (F _(1, 51)_ = 0.2955, P=0.5891) or sex (F _(1, 51)_ = 0.2944, P=0.5898). In female mice, SCVS significantly decreased the firing rate of dopaminergic neurons (control female= 4.50±0.86 Hz, n=11; SCVS female=2.73±0.35 Hz, n=12, p=0.033, Bonferroni post-hoc test). In male mice, however, SCVS had no significant effect on the firing rate (control male= 2.73±0.30 Hz, n=17; SCVS male=3.99±0.39 Hz, n=15, p=0.086, Bonferroni post-hoc test). These results show that SCVS induces reductions in tonic dopaminergic firing rate specifically in female mice.

### SCVS increases spontaneous inhibitory synaptic transmission onto VTA dopamine neurons in both males and females

We next sought to identify factors that may contribute to this reduction in dopaminergic firing rate in females. VTA dopaminergic neurons exhibit pacemaker firing and spend their lives in a perilously depolarized state, and are thus susceptible to robust inhibition by activation of GABA_A_ synapses through both direct hyperpolarization and shunting inhibition^46–50^. We therefore hypothesized that the reductions in VTA dopaminergic neuron firing rate we observe in females may be due to changes in inhibitory tone. To test this, we performed recordings of GABA_A_ mediated IPSCs from dopaminergic neurons in acute slices from the VTA of pitx3-GFP mice 1-4 days following the conclusion of SCVS. We first analyzed spontaneous IPSCs, which are a combination of action potential driven and action potential independent synaptic events (Figure 2A). In female animals, as expected, sIPSC frequency was higher in cells recorded from stressed mice (Figure 2B-C, control female = 1.80 ± 0.22 Hz, n=11; SCVS female = 3.80 ± 0.56 Hz, n=11). Intriguingly, we also found increased sIPSC frequency in dopaminergic neurons recorded from stressed male mice (Figure 2B-C, control male = 1.57±0.26 Hz, n=11; SCVS male = 3.45 ± 0.66 Hz, n=10). We found a significant effect of stress on sIPSC frequency (F _(1, 39)_ = 17.92, P=0.0001) but no significant effect of sex (F _(1, 39)_ = 0.4006, P=0.5305), or interaction between stress and sex (F _(1, 39)_ = 0.01321, P=0.9091). sIPSC amplitudes were unchanged following stress in both male and female mice (Figure 2D-E, control female = 33.89±3.79 pA, n=11; SCVS female = 42.23 ± 11.84 pA, n=11; control male 31.31±3.08 pA, n=11; SCVS male =33.95±3.442 pA, n=10), and there was no significant effect of stress (F_(1, 38)_ = 0.7348, P=0.3967), sex (F_(1, 38)_ = 0.7180, P=0.4021), or interaction between sex and stress (F_(1, 38)_ = 0.1980, P=0.6588). Taken together, this shows that SCVS increases presynaptic, but not postsynaptic, inhibitory tone onto dopaminergic neurons in both male and female mice.

**Figure 2:**
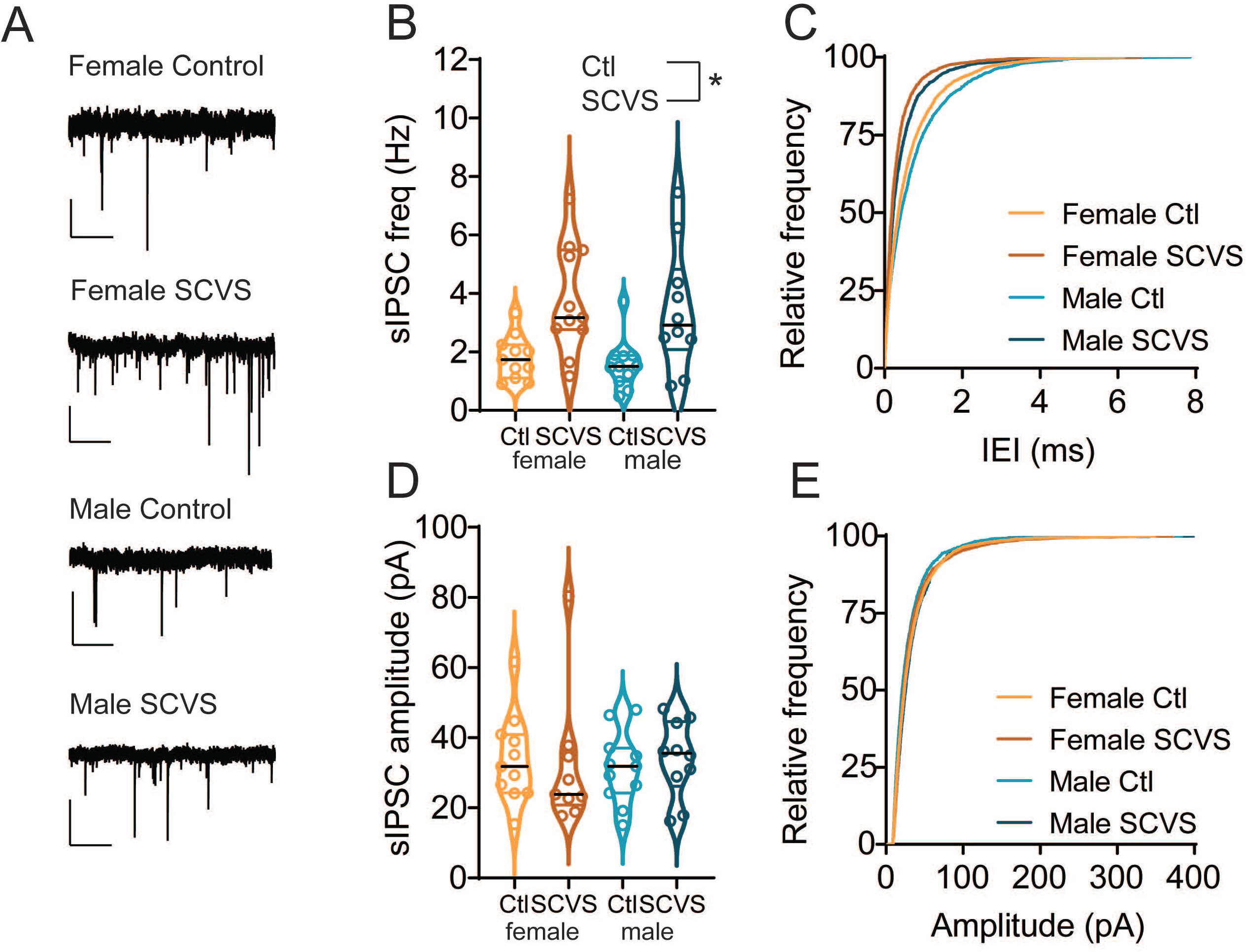
sIPSC frequency onto dopaminergic neurons is increased following SCVS in both male and female mice. A. Representative traces of spontaneous IPSCs from dopaminergic neurons. B. Average sIPSC frequency from VTA dopaminergic neurons from control and SCVS mice. *p<0.05, 2-way ANOVA, main effect of stress. C. Cumulative distribution of sIPSC inter-event intervals. D. Average sIPSC amplitudes from VTA dopaminergic neurons from control and SCVS mice. E. Cumulative distribution of sIPSC amplitudes. n=10-11/group. Scale bars=50 pA, 1 s.

Increases in sIPSC frequency can be caused by either an increase in release probability at presynaptic synapses or an increase in the firing rate of local cells that are active in the slice preparation. To distinguish between these possibilities, we collected miniature IPSCs in the presence of tetrodotoxin, which would eliminate any increase due to action-potential dependent release (Figure 3A). In contrast to the stress-induced increase in sIPSC frequency, mIPSC frequency was unchanged following stress (Figure 3B-C, control female = 1.91±0.22 Hz, n=11; SCVS female = 2.14 ± 0.22 Hz, n=10; control male 1.5±0.27 Hz, n=10; SCVS male =2.30±0.36 Hz, n=9), and there was no significant effect of stress (F_(1, 36)_ = 3.609, P=0.07), sex (F_(1, 36)_ = 0.2012, P=0.66), or interaction between sex and stress (F_(1, 36)_ = 1.05, P=0.3120). We again saw no stress-induced change in mIPSC amplitudes (Figure 3D-E, control female = 30.87±2.08 pA, n=11; SCVS female = 36.26 ± 5.56 pA, n=10; control male 31.50±3.72 pA, n=10; SCVS male =34.79±3.69 pA, n=9), and there was no significant effect of stress (F_(1, 36)_ = 1.25, P=0.27), sex (F_(1, 36)_ = 0.01, P=0.91), or interaction between sex and stress (F_(1, 36)_ = 0.07, P=0.79). Taken together, these data show that following SCVS, both male and female mice have an increase in inhibitory synaptic tone that is action potential dependent.

**Figure 3:**
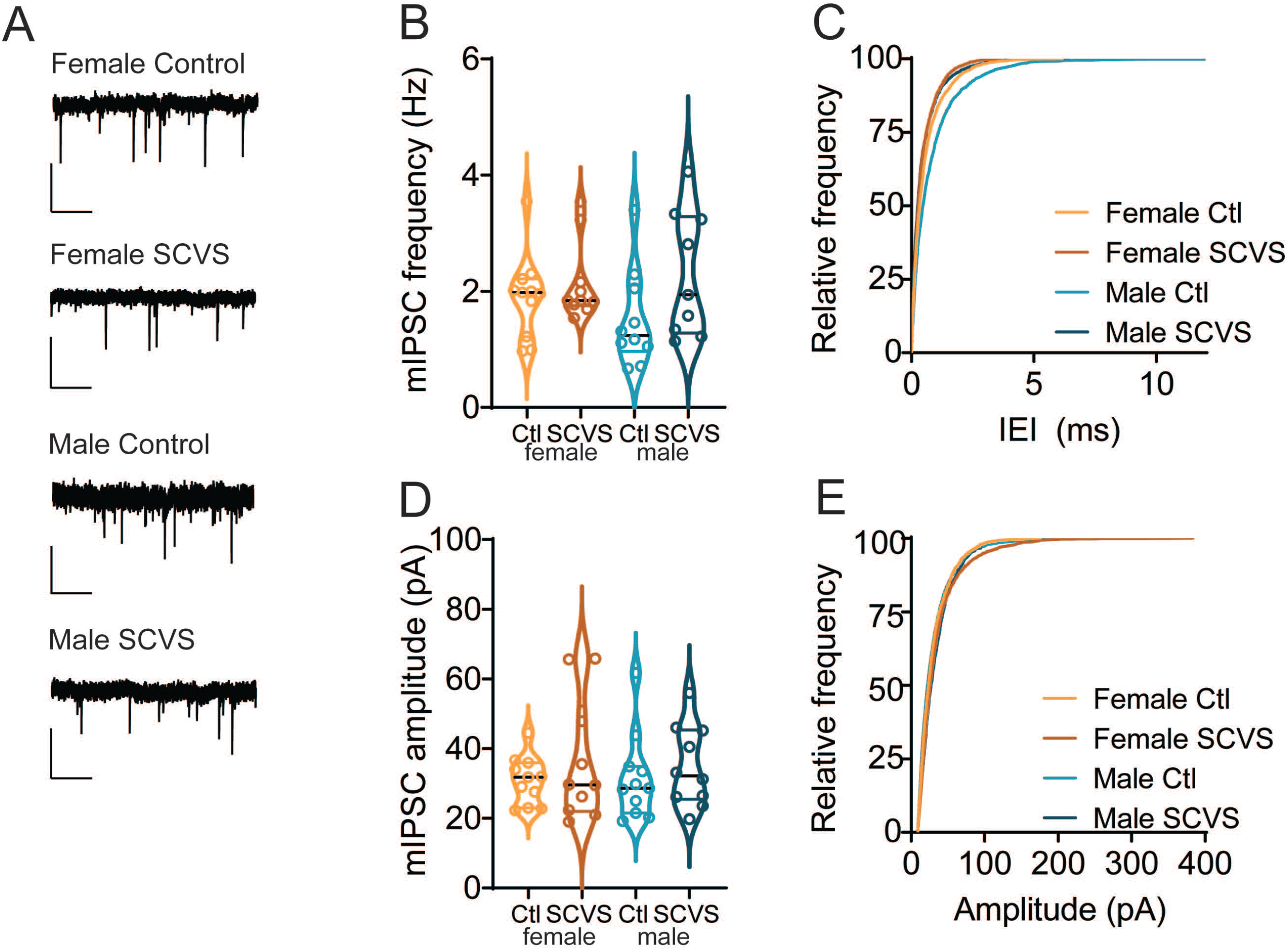
Frequency and amplitude of mIPSCs onto dopaminergic neurons are unchanged following SCVS. A. Representative traces of miniature IPSCs from dopaminergic neurons. B. Average mIPSC frequency from VTA dopaminergic neurons from control and SCVS mice. C. Cumulative distribution of mIPSC inter-event intervals. D. Average mIPSC amplitudes from VTA dopaminergic neurons from control and SCVS mice. E. Cumulative distribution of mIPSC amplitudes. n=9-11/group. Scale bars=50 pA, 1 s.

### SCVS increases firing rate of VTA GABA neurons in both sexes

VTA dopaminergic neurons receive GABA_A_ input from an array of sources, including both local GABAergic neurons and distal inputs such as the bed nucleus of the stria terminalis, ventral pallidum, rostromedial tegmental nucleus, and lateral hypothalamus^51^. Many of these sites are known to be sensitive to acute stress and could be altered by SCVS. However, given that our observed increase in inhibitory tone is preserved in an *ex vivo* slice, the source of this increase in inhibition must be cells that are present and connected to dopaminergic neurons in the slice. We therefore hypothesized that VTA GABAergic neurons might be the source of increased inhibitory tone following SCVS. To investigate this possibility, we performed SCVS in male and female Vgat-Cre:Ai14 mice that expressed tdTomato in GABAergic neurons and then performed cell-attached recordings to determine the basal firing rate of these cells. We found that the firing rate of Vgat+ VTA neurons was elevated following SCVS in both female (Figure 4A-B, control female 6.52 Hz ± 1.92 Hz, n=8; SCVS female 10.75 ± 1.72 Hz) and male (Figure 4A-B, control male 5.85 ± 1.42 Hz, n=7; SCVS male 12.74 ± 3.79 Hz, n=11). We found a significant effect of stress (F_1,34_=4.202, P=0.048) but no effect of sex (F_1,34_=0.059, P=0.809) or interaction between sex and stress (F_1,34_=0.241, P=0.627).

**Figure 4:**
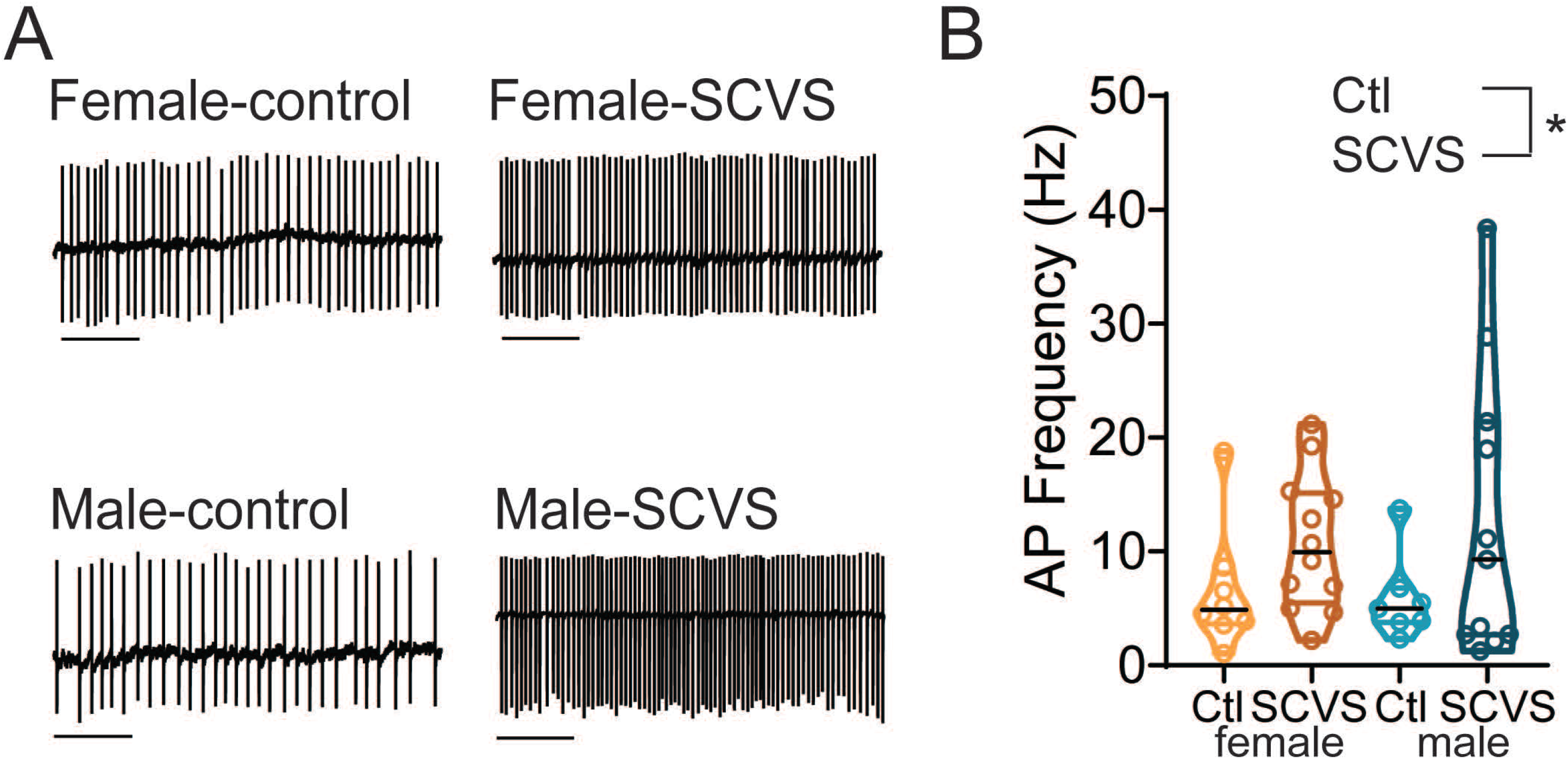
Firing rate of VTA GABA neurons is elevated after SCVS. A. Representative cell-attached traces from VTA GABA neurons. B. Average firing rate in control and SCVS mice. * P<0.05 main effect of stress, 2-way ANOVA. Scale bars=1s.

### Compensatory upregulation of excitatory transmission onto VTA dopamine neurons in males

Despite the SCVS-induced increase in local inhibitory tone onto dopaminergic neurons in both male and female mice, only female mice experience a decrease in the firing rate of dopaminergic neurons. This raises the possibility that there are homeostatic mechanisms in males that maintain the firing rate of dopaminergic neurons in the face of stress. With this in mind, we next investigated excitatory transmission onto VTA dopaminergic neurons in male and female pitx-GFP mice (Figure 5). Following six days of SCVS, we prepared acute VTA slices and performed whole cell recordings of spontaneous and miniature EPSCs. We first examined spontaneous EPSC frequency (Figure 5A-C). We found a significant interaction between stress and sex (F_1,40_=12.37, P=0.001), as well as a significant effect of stress (F_1,40_=4.560, P=0.039). There was no main effect of sex (F_1,40_=2.799, P=0.102). In female mice sEPSC frequency was unchanged following SCVS (control female 2.89 ± 1.04 Hz, n=9; SCVS female 1.10 ± 0.22 Hz, n=12, P=0.66, Bonferroni’s post-hoc test). However, in male mice we observed a significant increase in sEPSC frequency following six days of SCVS (control male 1.23 ± 0.30 Hz., n=14; SCVS male 6.31 ± 1.84 Hz, n=8; P=0.0006, Bonferroni’s post-hoc test).

**Figure 5:**
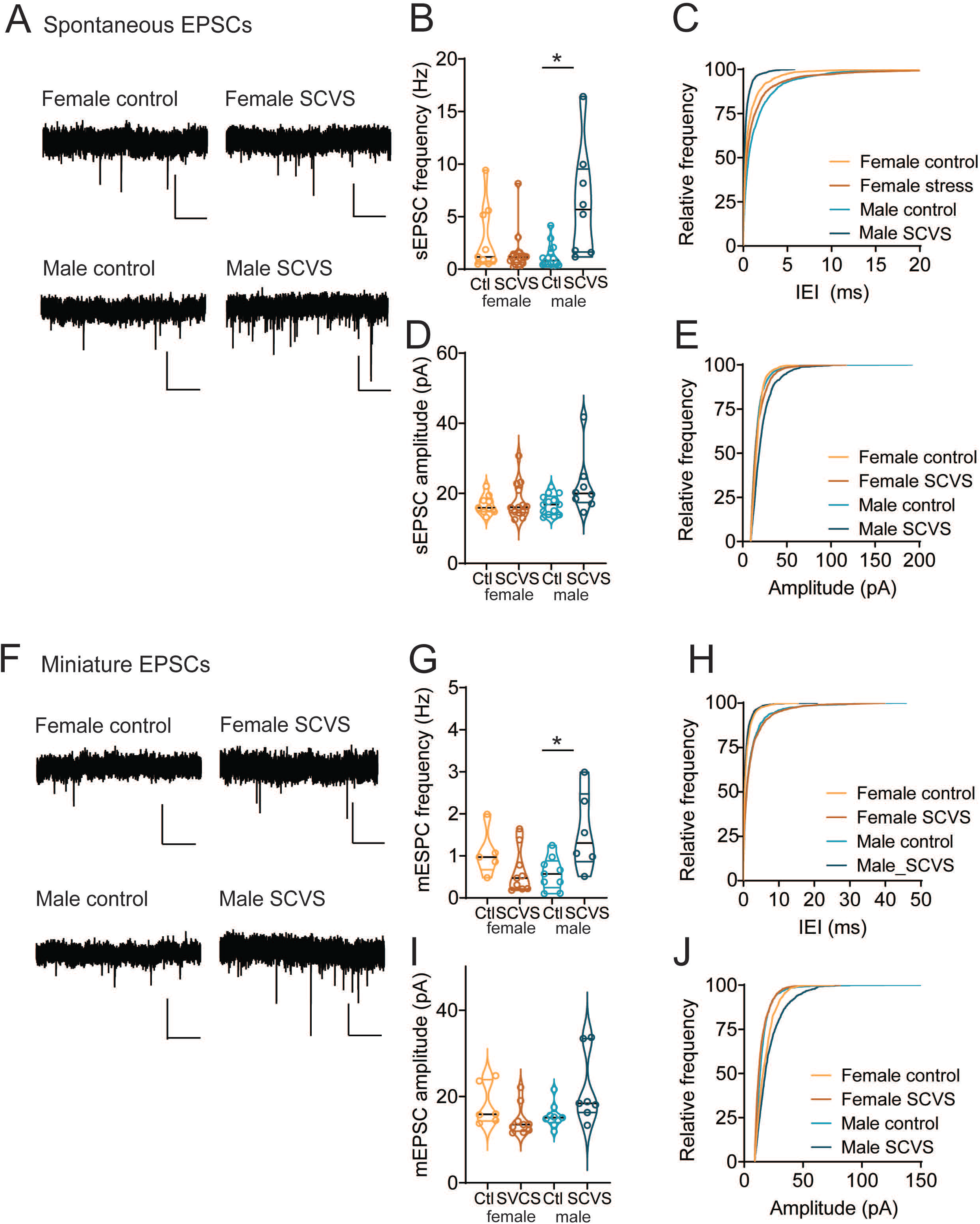
Increased frequency of excitatory transmission onto dopaminergic neurons in male mice following SCVS. A. Representative sEPSCs from VTA dopamine neurons. B. Average sEPSC frequency from VTA dopaminergic neurons from control and SCVS mice. *p<0.05, 2-way ANOVA followed by Bonferroni test. C. Cumulative distribution of sEPSC inter-event intervals. D. Average sEPSC amplitudes from VTA dopaminergic neurons from control and SCVS mice. E. Cumulative distribution of sEPSC amplitudes. F. Representative mEPSCs from VTA dopamine neurons. G. Average mEPSC frequency from VTA dopaminergic neurons from control and SCVS mice. *p<0.05, 2-way ANOVA followed by Bonferroni test. H. Cumulative distribution of sEPSC inter-event intervals. I. Average sEPSC amplitudes from VTA dopaminergic neurons from control and SCVS mice. J. Cumulative distribution of sEPSC amplitudes n=8-12/group. Scale bars=20 pA, 1 s.

**Figure 6:**
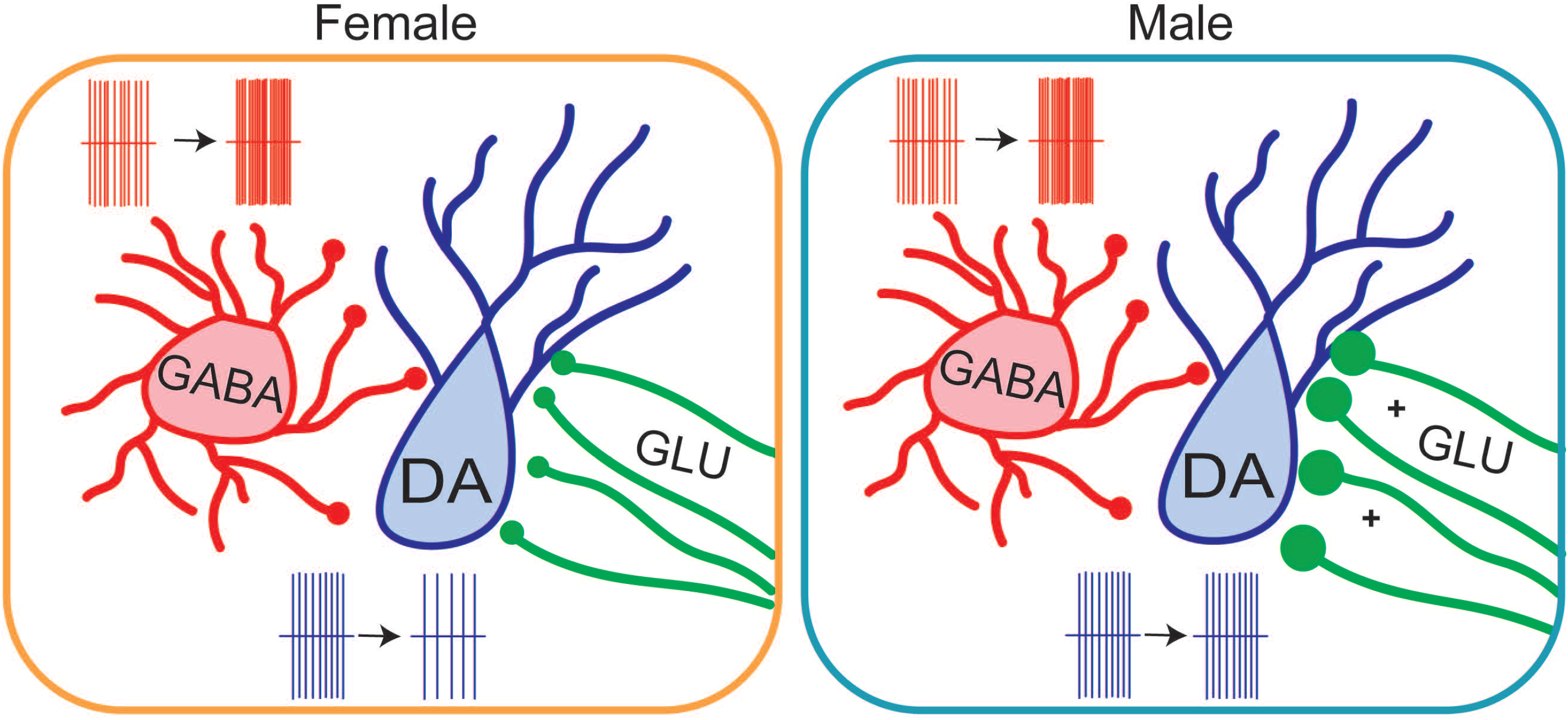
VTA adaptations following SCVS in female and male mice. In both female and male mice, SCVS leads to an increase in firing of VTA GABA neurons and an increase in inhibitory tone onto dopaminergic neurons. In female mice, this is accompanied by a decrease in the tonic firing rate of dopaminergic neurons. In male mice, however, upregulation of excitatory synapses leads to a maintenance of dopaminergic firing rate.

In contrast, no effect of sex (F_1,40_=0.46, P=0.50) or stress (F_1,40_=2.52, P=0.12), or interaction between the two (F_1,40_=0.56, P=0.46) was seen when looking at sEPSC amplitude (Figure 5D-E, control female = 16.81±0.91 pA, n=9; SCVS female = 17.82 ± 1.44 pA, n=13; control male 16.73±0.75 pA, n=14; SCVS male =19.52±1.25 pA, n=8).

To examine EPSC properties independently of action potentials, we also collected mEPSCs following SCVS (Figure 5F). As with sEPSCs, we found a significant interaction between stress and sex on mEPSC frequency (Figure 5G-H, F_1,25_=9.582, P=0.005), but no independent effect of stress (F_1,25_=1.408, P=0.257) or sex (F_1,25_=0.894, P=0.354). Dopaminergic neurons from male mice showed a significant increase in mEPSC frequency following SCVS (control male 0.58 ± 0.13 Hz., n=9; SCVS male 1.57 ± 0.38 Hz, n=6; P=0.009, Bonferroni’s post-hoc test), while no significant change was seen in females (control female 1.07 ± 0.25 Hz, n=5; SCVS female 0.63 ± 0.18 Hz, n=9, P=0.40, Bonferroni’s post-hoc test). Surprisingly, we saw a significant interaction between sex and stress on mEPSC amplitudes (Figure 5I-J, F_1,27_=6.5899, P=0.014). Dopaminergic neurons from male mice showed an increase in mEPSC amplitude following SCVS (control male 15.51 ± 0.92 pA., n=9; SCVS male 21.71 ± 3.14 pA, n=7; P=0.045, Bonferroni’s post-hoc test). No change was seen in mEPSC amplitudes from female mice (control female 18.06 ± 1.97 pA., n=6; SCVS female 14.54 ± 1.20 pA, n=9; P=0.399, Bonferroni’s post-hoc test). We again saw no significant effect of stress (F_1,27_=0.5258, P=0.475) or sex (F_1,27_=0.1.553, P=0.223) on mEPSC amplitude. Together, this shows that following stress, male mice exhibit a robust upregulation of excitatory synapses.

## Discussion

### Sex-specific alterations in tonic dopaminergic firing following subchronic stress

In this study, we used subchronic variable stress (SCVS) to investigate regulation of the ventral tegmental area circuitry in a model of female-specific vulnerability to stress. We found that female, but not male mice, showed a decrease in the *ex vivo* firing rate of dopaminergic neurons within the ventral tegmental area following stress. A prior study has shown no effect of SCVS on *ex vivo* firing rate of putative dopaminergic neurons^52^, however significant differences in selection criteria for dopaminergic neurons may underlie this difference. While we did not specifically label neurons by projection target in this study, our selection criteria (GFP expression in Pitx3-GFP mice, lateral location and I_h_+) are consistent with recording exclusively dopaminergic cells and biasing our population towards those that project to the lateral nucleus accumbens shell^37–39^. This reduction in firing is therefore likely to lead to reduced tonic levels of dopamine in the accumbens and potentially other projection targets.

SCVS leads to several pronounced behavioral changes specifically in female mice, including reduced sucrose preference^26, 30^, decreased social interaction^28^, increased avoidance behavior in the novelty-suppressed feeding assay^26, 53^, reduced grooming^26, 53^, and increased passive coping in the forced swim test^27, 53^. Stress-induced changes in these behaviors have been closely linked to activity of dopaminergic neurons. Behaviors such as sucrose preference^21^, sociability^44^ and avoidance behavior^43^ can be altered by manipulation of the dopaminergic system. Thus, our finding that SCVS reduces tonic dopaminergic firing specifically in female mice is consistent with SCVS-induced female-specific alterations in dopamine-dependent behavior^26, 30, 53^.

It is important to note that the relationship between chronic stress and VTA dopaminergic function is a complicated one. While chronic unpredictable stress dampens activity of dopaminergic neurons and reduces dopamine release in the nucleus accumbens^21, 54–58^, other models, such as chronic social defeat stress (CSDS) increase activity of these same dopaminergic neurons^15, 16^, and silencing dopaminergic neurons reverses social avoidance and sucrose preference deficits induced by CSDS^15^. There are several explanations for these contrasting results. For one example, the relationship between activity in the VTA-NAc dopaminergic neurons and social and sucrose reward could be an inverted U-shaped curve, where too little or too much dopaminergic activity has similar effects on behavior. This dose response relationship has been proposed to underlie dopaminergic effects on cognition in the prefrontal cortex^59^. Differences in post-stress configuration of the postsynaptic circuit in the nucleus accumbens that alter the “optimal” set point for dopaminergic tone following stress may also underlie these effects. Prior work has suggested that the duration of stress may underlie differential roles of dopaminergic neurons in post-stress sequelae, as CSDS protocols typically last 10 days and chronic unpredictable stress protocols often extend for at least four weeks. However, our results here show that at least in female mice, decreases in dopaminergic firing are possible following only six days of stress.

### SCVS increases inhibitory tone onto DA neurons in both males and females

Even though only female mice exhibited SCVS-induced decreases in the firing rate of VTA dopaminergic neurons, inhibitory tone onto these neurons was increased in *both* male and female mice. Spontaneous, but not miniature, IPSC frequency onto dopaminergic neurons was increased, indicating that the increase was mediated by action potential dependent release rather than changes in presynaptic release probability or the number of release sites. This suggests that this increase in inhibition arises from a local source; consistent with this, we see an increase in the firing rate of VTA GABA neurons in both male and female mice following SCVS. It is notable that there is considerable variability among firing rates within VTA GABA neurons following SCVS, suggesting that increases in firing may not be uniform across all GABA neuron subtypes. VTA GABA neurons are known to be heterogeneous^60^, but the nature of this heterogeneity remains enigmatic. There are no consensus markers for genetic access to distinct subclasses of cells, or even of what subclasses exist; we therefore used a VGAT-Cre line to label GABAergic neurons for this study. Our recordings are potentially arise from a mix of interneurons that only connect locally and projection neurons that send axons to distal targets and may or may not have local collaterals contacting other cells within the VTA^60^. While we have focused on local effects within the VTA in this study, we cannot rule out that there is also increased GABAergic tone in distal targets of VTA GABA neurons. These projections are important regulators of reward and avoidance behavior independently of their effects on dopaminergic neurons^60–63^, and upregulation of their activity is likely to also have significant consequences for behavior.

VTA GABA neurons are robustly activated by acute stressors^49, 64, 65^, here we demonstrate that longer-term stressors can also lead to persistent changes in firing of these cells. VTA GABA neurons receive of wealth of stress-sensitive excitatory and inhibitory inputs from regions such as the prefrontal cortex, bed nucleus of the stria terminalis, hypothalamus, and dorsal raphe nucleus^60, 66, 67^. Alterations in the strength of any of these inputs, or the balance between them, could drastically change the firing rate of these cells. Furthermore, VTA GABA neurons are influenced by a number of stress-related peptides and neuromodulators including opioids^34, 68^, CRF^69^, NPY^69^, serotonin^70^, and acetylcholine^70, 71^. Alternatively, persistent increases in activity of these neurons could be due to intrinsic factors regulating excitability, or interactions between intrinsic and extrinsic plasticity. For example, repeated restraint stress alters the chloride reversal potential in VTA GABA neurons, shifting GABAergic inputs from inhibitory to excitatory and enhancing inhibition of VTA dopaminergic neurons^72^. Future studies investigating mechanisms of regulation of these cells during and following SCVS will be valuable.

### Compensatory upregulation of excitatory synapses in males

Despite the increase in inhibitory tone onto dopaminergic neurons in both male and female mice, only females show a decrease in the dopaminergic neuron firing rate. Intriguingly, we find that males exhibit a strong upregulation of excitatory tone onto dopaminergic neurons. VTA dopamine neurons receive a number of excitatory inputs from both local glutamatergic neurons^73, 74^ and distal sources such as the LH, BNST, PFC, and DRN^51^. This increase is not sensitive to tetrodotoxin, indicating that is likely mediated at the synaptic level by an increase in release probability and/or sprouting of new release sites. Thus, the brains of male, but not female, mice appear to compensate for a stress-induced increase in inhibition with a counterbalancing increase in excitation.

It remains unclear what drives this increase in excitatory signaling. Androgen hormones such as testosterone are one factor that could contribute to male-specific changes. Androgen-dependent alterations in the excitability of ventral hippocampal neurons contribute to male resistance to SCVS^30^. Similar androgen dependence is possible in the VTA as both gonadal and locally produced androgen hormones and androgen receptors are present in the VTA^75–77^. Further investigation of the timing of this adaptation will also be revealing. If increases in the firing rate of VTA GABA neurons precede upregulation of excitatory synapses, it suggests that the change in excitatory synapses could be a homeostatic response to an increase in inhibition. Several studies have shown that pharmacological manipulation of GABA_A_ receptor function or GABAergic tone in the VTA can trigger adaptations in excitatory synapses onto dopaminergic neurons^78, 79^; whether a similar mechanism exists following stress remains to be seen. Intriguingly, while six days of SCVS is not sufficient to induce behavioral changes in male mice, repetition of this for twenty-one or more days leads to anhedonia and avoidance in both female and male mice^28, 53, 80^. Whether this later timepoint is associated with change in dopaminergic function in males, and whether this requires reversal of the increase in excitatory synapses induced early in stress will be an important topic for future investigations.

These data are in line with a wealth of studies showing that both vulnerability and resistance to stress are active processes^81^. Just as maladaptive behavioral and physiological changes require alterations in genes, molecules, circuits, and systems in the brain, maintaining or regaining behavioral homeostasis in face of environmental challenges requires constant adaptation^82^. Whether a behavioral or physiological change is adaptive or maladaptive is context dependent, and changes that are adaptive in one context may be detrimental in another. In male mice, an upregulation of excitatory synapses appears to protect against loss of dopaminergic tone. However, excitatory synapses are major driver of phasic bursting in VTA dopaminergic neurons^83–87^. Thus, an adaptation that protects against diminished tonic dopaminergic signaling could prime the mesolimbic system for enhanced phasic dopaminergic signaling, inducing a latent vulnerability to heightened response to motivational stimuli and aberrant reinforcement learning in male mice. Taken together, our data reveal that subchronic variable stress induces both shared and divergent mechanisms of regulation of VTA dopaminergic neurons. These findings have significant implications for post-stress regulation of the mesolimbic dopaminergic circuitry, and future investigation of the mechanisms of these changes may provide pathways for the development of sex-specific treatments for stress-linked neuropsychiatric illness.

## Acknowledgements

The authors thank Dr. Paul Marvar for use of footshock boxes, Dr. Zhe Yu for advice on immunohistochemistry, and Dr. Kevin Wickman for the Pitx3-GFP mice. This work was performed in part at the George Washington University Nanofabrication and Imaging Center (GWNIC).

## Author Contributions

CB and AMP designed the research. CB, AKisner, and AMP collected and analyzed electrophysiology data. BVT, AKhurana, and CRC performed stress experiments. MW performed immunohistochemistry and collected microscope images. CB and AMP contributed to data interpretation. AMP wrote the paper with input from the other authors.

